# A comparative study of the capacity of mesenchymal stromal cell lines to form spheroids

**DOI:** 10.1101/834390

**Authors:** Margaux Deynoux, Nicola Sunter, Elfi Ducrocq, Hassan Dakik, Roseline Guibon, Julien Burlaud-Gaillard, Lucie Brisson, Louis-Romée le Nail, Olivier Hérault, Jorge Domenech, Philippe Roingeard, Gaëlle Fromont, Frédéric Mazurier

## Abstract

Mesenchymal stem cells (MSCs)-derived spheroid models favor maintenance of stemness, *ex vivo* expansion and transplantation efficacy. Spheroids may also be considered as useful surrogate models of the hematopoietic niche. However, accessibility to primary cells, from bone marrow (BM) or adipose tissues, may limit their experimental use and the lack of consistency in methods to form spheroids may affect data interpretation. In this study, we aimed to create a simple model by examining the ability of cell lines, from human (HS-27a and HS-5) and murine (MS-5) BM origins, to form spheroids, compared to primary human MSCs (hMSCs). Our protocol efficiently allowed the spheroid formation from all cell types within 24 hours. Whilst hMSCs-derived spheroids began to shrink after twenty-four hours, the size of spheroids derived from cell lines remained constant during three weeks. The difference was partially explained by the balance between proliferation and cell death, which could be triggered by hypoxia and induced oxidative stress. Our results demonstrate that, unlike hMSCs, MSC cell lines make reproductible spheroids that are easily handled. Thus, this model could help in understanding mechanisms involved in MSC functions and may provide a simple model by which to study cell interactions in the BM niche.

## Introduction

Over the last two decades, extensive studies have attempted to characterize mesenchymal stem cell (MSC). Initially described in the bone marrow (BM), MSCs were later found in almost all adult and fetal tissues [1]. Their classification rapidly suffered from a lack of clear phenotypical definition. Therefore, in 2006, the International Society for Cellular Therapy (ISCT) defined MSCs according to three minimal criteria: adherence to plastic, specific cell surface markers and multipotent potential. Indeed, MSCs are classically described as stem cells that are able to differentiate into osteoblasts, adipocytes and chondroblasts [2], making them an attractive source of cells in regenerative medicine. Subsequent studies have also established their ability to differentiate into cardiomyocytes [3], neurons [4], epithelial cells [5] and hepatocytes [6]. The discovery of the multiple functions of MSC, such as those involved in the anti-inflammatory response [7] and in injury repair [8,9] confirmed them as promising cellular tools in regenerative medicine.

Furthermore, MSCs represent a key component of the BM microenvironment supporting normal hematopoiesis through the regulation of stem cell renewal and differentiation processes, but also fueling malignant cells and protecting them from therapeutic agents [10]. As such, primary MSCs have often been used as feeder layers in long-term co-culture of hematopoietic cells *in vitro* in preclinical studies [11]. With the aim of standardization, the murine MS-5 cell line became the gold-standard for both normal or malignant hematopoietic cell culture [12]. This robust co-culture model has been widely used and has contributed to the characterization of hematopoietic stem cells (HSC) [11]. This 2D system, while closer to BM physiology than the culture of hematopoietic cells alone, still lacks the three-dimensional complexity of the BM niche. Thus, although widely used, it is certainly not sufficiently consistent at predicting *in vivo* responses [13]. Therefore, a 3D system might be a better alternative to mimic the BM microenvironment.

Critically, the culture leads to rapid loss of MSC pluripotency and supportive functions. Therefore, a wide range of techniques to form 3D MSC structures, from the simplest spheroids to the more complex matrix-based structures, have been proposed [14]. Studies of spheroids, also called mesenspheres, were mostly dedicated to the examination of MSC stemness and differentiation abilities, such as osteogenesis, in order to improve their *in vitro* expansion and transplantation efficacy in regenerative medicine [15,16]. Furthermore, this model has also been tested as a surrogate niche for hematopoietic cells [17–23]. Spheroids take advantage of the ability of MSCs to self-aggregate, which is improved by using various approaches such as low adhesion plates, natural and artificial (centrifugation) gravity, cell matrix or more complex scaffolds [13,14,24,25]. Classically, studies have used human primary MSCs, from BM, cord blood and lipoaspirate, or rodent sources [15,26].

Although immortalized MSCs, or well characterized cell lines, could bypass the lack of primary cells and avoid the variability involved with use of primary human MSCs (hMSCs) samples, they are rarely employed to make spheroids [27,28]. Cell lines would also allow better standardization of the spheroid formation protocol. In this study, we examined the spheroid-forming capacity of two human cell lines (HS-27a and HS-5) and the murine gold-standard MS-5, in comparison with hMSCs. We defined a simple and fast method using standard matrix to form spheroids and characterized them in terms of physical features, cell proliferation and death.

## Materials and methods

### Cell culture and reagents

Murine MS-5 bone marrow (BM) stromal cell line was kindly provided by Mori KJ (Niigata University, Japan) [29]. Human HS-27a and HS-5 BM stromal cell lines were purchased from the American Type Culture Collection (ATCC CRL-2496 and CRL-11882, respectively). Primary human MSCs (hMSCs) were obtained by iliac crest aspiration from informed consent patients undergoing orthopedic surgery (Cardiovascular Surgery Department, Trousseau Hospital, Tours, France). HS-27a and HS-5 cell lines were cultured in RPMI 1640 (Life Technologies, Villebon-sur-Yvette, France) and hMSCs and MS-5 in MEM Alpha (Life Technologies). All medium were supplemented with 10 % heat-inactivated fetal bovine serum (FBS), 2 mM L-glutamine, 100 U/mL penicillin and 100 μg/mL streptomycin (all from Life Technologies). For hMSCs culture only, 0.004 % of recombinant human FGF basic (FGF-2, R&D Systems, Abingdon, United Kingdom) was added. Cells were maintained in a saturated humidified atmosphere at 37°C and 5 % CO_2_. HS-27a, HS-5 and MS-5 cell lines were used for experiments between passages 5 and 20, and hMSCs at passage 2.

### Spheroids formation

For one spheroid, 30,000 cells were cultured in 100 μL of medium, supplemented by 0.25 % to 1 % of either Methocult™ SF H4236 or H4100 (StemCell, Grenoble, France), and seeded in U-bottomed 96-well plate (Sarstedt, Marnay, France). The medium was the same as that of the normal culture for each cell line but supplemented with heat inactivated FBS to reach 15 %. At days as detailed, microscopic analysis was performed using a Leica DMIL microscope (Leica, Nanterre, France), coupled to a DXM1200F camera (Nikon, Champigny-sur-Marne, France). To determine the number of cells in each spheroid over time, 12 spheroids per experiment were pooled and dissociated with 2 mg/mL collagenase 1A (Sigma-Aldrich, Saint-Quentin-Fallavier, France), 10 min at 37°C, with agitation every two minutes, and then counted by the trypan blue exclusion assay.

### Time-lapse video

Automatic acquisitions were performed on a Nikon Eclipse TI-S microscope, coupled to a DS Qi2 camera (Nikon). The system includes a cage incubator (Okolab, Pozzuoli, NA, Italy) controlling temperature and level of CO_2_. Analyses were performed using both NIS Element BR (Nikon) and Fiji/ImageJ softwares.

### Scanning electron microscopy

Spheroids were fixed by incubation for 24 h in 4 % paraformaldehyde, 1 % glutaraldehyde in 0.1 M phosphate buffer (pH 7.2). Samples were then washed in phosphate-buffered saline (PBS) and post-fixed by incubation with 2 % osmium tetroxide for 1 h. Spheroids were then fully dehydrated in a graded series of ethanol solutions, and dried in hexamethyldisilazane (HMDS, Sigma-Aldrich). Finally, samples were coated with 40 Å platinum, using a PECS 682 apparatus (Gatan, Evry, France), before observation under an Ultra plus FEG-SEM scanning electron microscope (Zeiss, Marly-le-Roi, France).

### Transmission electron microscopy

Spheroids were fixed by incubation for 24 h in 4 % paraformaldehyde, 1 % glutaraldehyde in 0.1 M phosphate buffer (pH 7.2). Samples were then washed in phosphate-buffered saline (PBS) and post-fixed by incubation with 2 % osmium tetroxide for 1 h. Spheroids were then fully dehydrated in a graded series of ethanol solutions and propylene oxide. Impregnation step was performed with a mixture of (1:1) propylene oxide/Epon resin, and then left overnight in pure resin. Samples were then embedded in Epon resin, which was allowed to polymerize for 48 h at 60°C. Ultra-thin sections (90 nm) were obtained with an EM UC7 ultramicrotome (Leica). Sections were stained with 5 % uranyl acetate (Agar Scientific, Stansted, United Kingdom), 5 % lead citrate (Sigma-Aldrich) and observations were made with a transmission electron microscope (Jeol, JEM 1011, Croissy-sur-Seine, France).

### Immunohistochemistry

At least five spheroids per conditions were pooled, fixed in formalin, embedded in paraffin and cut in 3-4 μm sections on Superfrost Plus slides. Slides were deparaffinized, rehydrated and heated in citrate buffer pH 6 for antigenic retrieval. After blocking for endogenous peroxidase with 3 % hydrogen peroxide, the primary antibodies were incubated. The panel of primary antibody included anti-HIF-1α (Abcam ab51608, Paris, France) (dilution 1/200, incubation 1 h), VEGF-A (Abcam ab1316, dilution 1/200, incubation 1 h), HO-1 (Abcam ab52947, dilution 1/1 000, incubation 1 h), CA-IX (Novocastra clone TH22, Nanterre, France) (dilution 1/100, incubation 20 min), and Ki-67 (DakoCytomation clone 39-9, Glostrup, Denmark) (dilution 1/50, incubation 30 min). Immunohistochemistry was performed with either the automated BenchMark XT slide stainer (Ventana Medical System Inc.) using OptiView Detection Kit (Ventana Medical System Inc.) (for CA-IX and Ki-67), or manually using the streptavidin-biotin-peroxidase method with diaminobenzidine as the chromogen (Kit LSAB, DakoCytomation). Slides were finally counterstained with haematoxylin. Negative controls were obtained after omission of the primary antibody or incubation with a non-specific antibody.

### Quantitative real-time PCR

Total RNAs were extracted using TRIzol reagent (15596-026, Life Technologies) and reverse transcription was performed with the SuperScript™ VILO™ cDNA Synthesis Kit (11754-050, Invitrogen, Villebon-sur-Yvette, France), both according to the manufacturer’s procedures. qRT-PCR was performed on a LightCycler® 480 (Roche, Switzerland) with the LightCycler® 480 Probes Master (04887301001, Roche). *GAPDH*, *ACTB*, *RPL13A* and *EF1A* genes were used as endogenous genes for normalization. Primer sequences (S1 Table) were designed with the ProbeFinder software (Roche), and all reactions were run in triplicate.

### Cell cycle analysis

Spheroids were dissociated with 2 mg/mL collagenase 1A (Sigma-Aldrich), 10 min, at 37°C, with agitation every two minutes. Cells were fixed with 2 % paraformaldehyde/0.03 % saponin for 15 min at room temperature (RT), and washed three times for 5 min with 10 % FBS/0.03 % saponin. Cells were then stained with 7-Aminoactinomycin D (7-AAD, Sigma-Aldrich) and an AF488-conjugated anti-KI-67 antibody (BD Biosciences, Le Pont de Claix, France) or the AF488-conjugated IgG_1_ isotype control (BD Biosciences). Experiments were performed on Accuri™ C6 flow cytometer (BD Biosciences) and data were analyzed with the FlowJo V10.4.1 software (Tree Star Inc.).

### Statistical analysis

All statistical analyses were performed using R software. The Mann-Whitney test was used to compare two conditions and Kruskall-Wallis for multiple comparisons, followed by a Dunn’s *post hoc* test. The threshold for significance was set up to a p-value of 0.05.

## Results

### Establishment of hMSC-derived spheroids by cell aggregation method

Among different methods to form MSC-spheroids, we followed an approach based on cell aggregation in methylcellulose-based medium [27] (Fig 1A). To establish a protocol that is simple, reproducible and compatible with hematopoietic cell culture, two commercial methylcelluloses commonly used for hematopoietic cell assays were tested. In general, a range from 0.01 to 1 % of methylcellulose was used [27,30–33], so we tested three different concentrations (0.25, 0.5 and 1 %). We also tested a hanging drop technique [31,33–35] and the previously described U-bottomed 96-well plates methods [27,30,32,33,36]. Both techniques worked well for primary hMSCs but the second was more appropriated for further analyses and offered lesser dehydration (data not shown). The SF H4236 methylcellulose at a concentration of 0.5 % was adopted because it generated one spheroid per well with lower condensation aspect for primary hMSCs (Fig 1B). Under these culture conditions, hMSCs were able to form spheroids rapidly, in as little as five hours of culture (S1 Video), which is consistent with previous studies [27,32,37].

**Fig 1.**
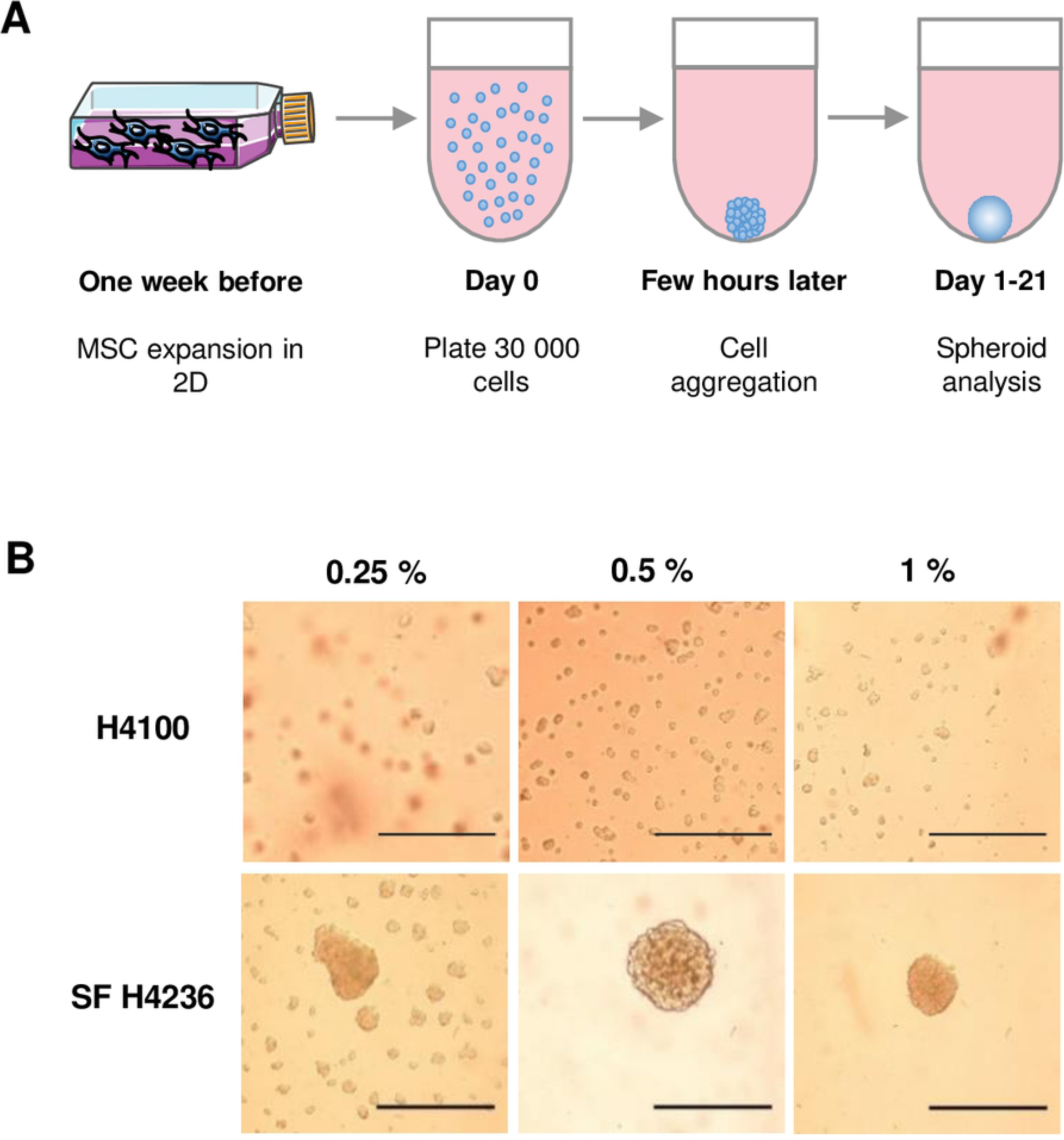
Spheroids formation from hMSCs. (A) Schematic representation of experimental plan. (B) 30,000 hMSCs per well were seeded into U-bottomed 96-well in medium containing 0.25 %, 0.5 % or 1 % of methylcellulose (Methocult™ H4100 or SF H4236). Microscopy analysis was performed after 24 h (scale bars = 500 μm).

### Formation of spheroids from MSC cell lines

The spheroid-forming capacity was followed for two human cell lines, HS-27a and HS-5, and compared to that of hMSCs. These cell lines have been obtained by immortalization of hMSCs with the papilloma virus E6/E7 genes [38,39]. HS-27a cells support hematopoietic stem cell maintenance (self-renewal, formation of cobblestone areas), whereas HS-5 cells mainly sustain proliferation and differentiation [38–40]. Unlike hMSCs, they retained the ability to form spheres but required about 10 hours to make rounded spheroids (S2 and S3 Videos). Although MSCs of various origins formed spheroids of equivalent sizes (about 300 μm of diameter) after 24 hours, hMSCs-derived spheroids rapidly condensed and reached half of their initial perimeter after 14 days of culture (Fig 2A and B). In contrast to hMSCs, the perimeter of spheroids resulting from both cell lines remained constant during three weeks. Knowing that hMSCs and cell lines may differ in their growth properties, we used the murine MS-5 cell line that has contact inhibition [29]. This cell line was able to quickly form spheroids similarly to the other cell lines (S4 Video). It is noteworthy that MS-5 cells initially formed a flat multilayer disk of cells prior to contracting into spheres. Similarly to the spheroids derived from human cell lines, spheroids from MS-5 cells kept the same size over time (S1A and S1B Fig). This suggests that shrinking might be an intrinsic property or extracellular matrix (ECM) composition of primary cells rather than related to cell proliferation control. We thus examined whether the difference in the size maintenance between various MSCs might be attributed to the cell number per spheroid. In order to quantify the viable cells, spheroids were dissociated at different timepoints after seeding. In accordance with the decrease in circumference, the number of cells per spheroid for hMSCs dramatically dropped within seven days (Fig 2C), which was in agreement with other studies [31,35]. Remarkably, although keeping the same size, HS-27a-derived spheroids, as well as the MS-5 ones, had lost viable cells similarly to hMSCs (Fig 2C and S1C Fig). In contrast, HS-5-derived spheroids had less obvious decrease in cell number with time (Fig 2C). Overall, the size reduction does not seem to be strictly attributable to reduced cell number in spheroids and could be possibly attributed to other factors such as the ECM composition.

**Fig 2.**
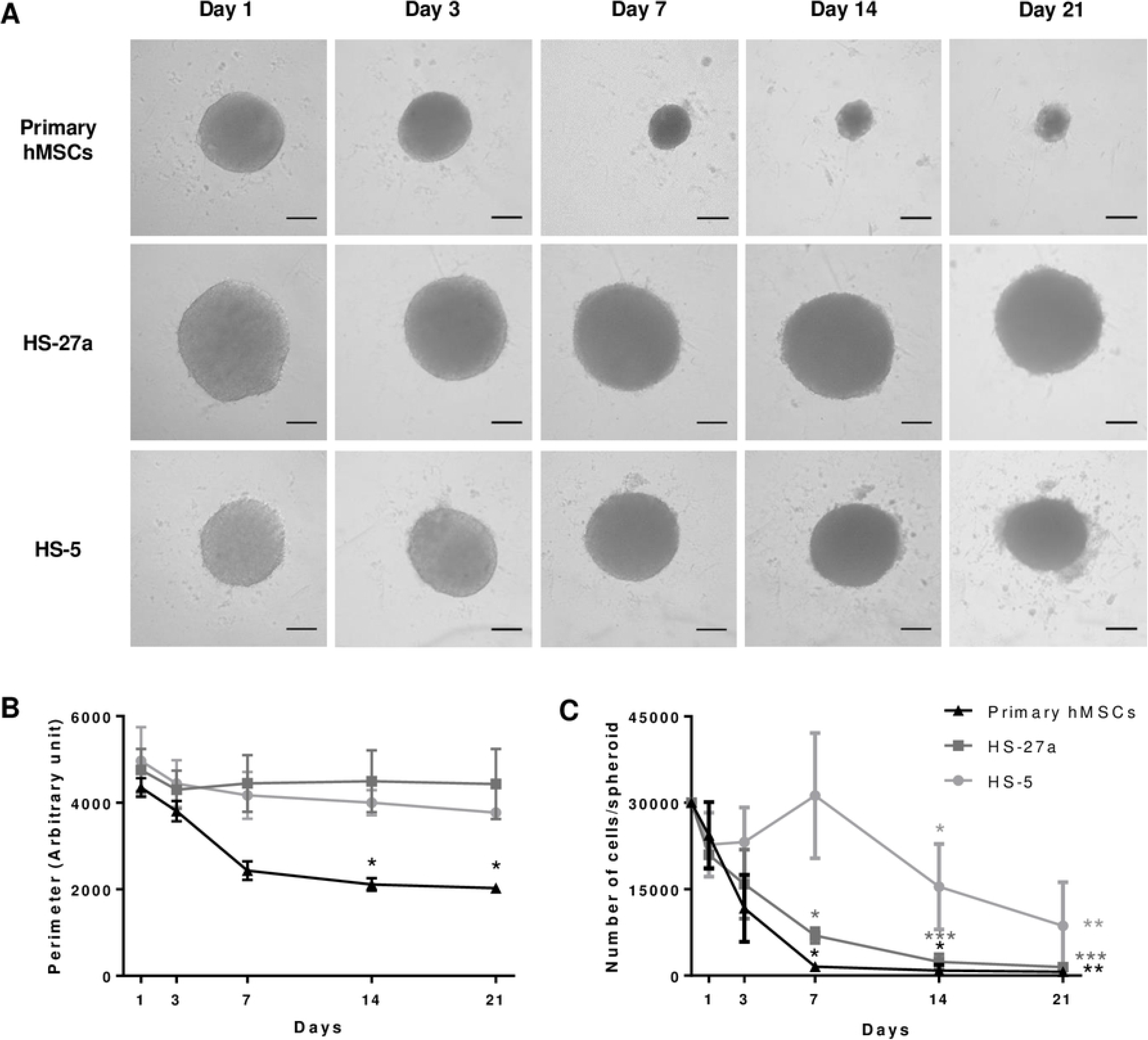
Follow up of the spheroids derived from various MSCs. (A) Microscopy analysis of hMSCs-, HS-27a- and HS-5-derived spheroids over 21 days in culture (scale bars = 100 μm). (B) Perimeter was measured with an arbitrary unit; each experiment is the mean of at least 10 spheroids from n = 3 experiments. Data are mean ± SD; * compared to day 1; * p ≤ 0.01. (C) Number of living cells per spheroids over 21 days in culture (hMCSs n = 3; HS-27a and HS-5 n = 4; each experiment is the mean of 12 spheroids).

### Electron microscopy observation of the MSCs-derived spheroids

Scanning electron microscopy (SEM) confirmed the shrinking of hMSC-derived spheroids (Fig 3A and S1D Fig). SEM also revealed at higher magnification that spheroids from hMSCs are highly cohesive, showing tight intercellular connections forming a flat surface, whereas HS-27a, HS-5 and MS-5 spheroids exhibited more rounded cells at their surface (Fig 3B and S1E Fig). This phenomenon intensified over the time and, in line with the assumption that ECM composition is different, may explain the size reduction of hMSC-derived spheroids compared to the cell lines. From day 7 for cell lines and day 14 for primaries, spheroid structure began to change, suggesting a progressive cell death. Further analysis by transmission electron microscopy (TEM) to investigate the ultrastructure of the cells within the spheroids showed the appearance of a progressive cell injury, thus confirming induced cell death (Fig 3C and S1F Fig).

**Fig 3.**
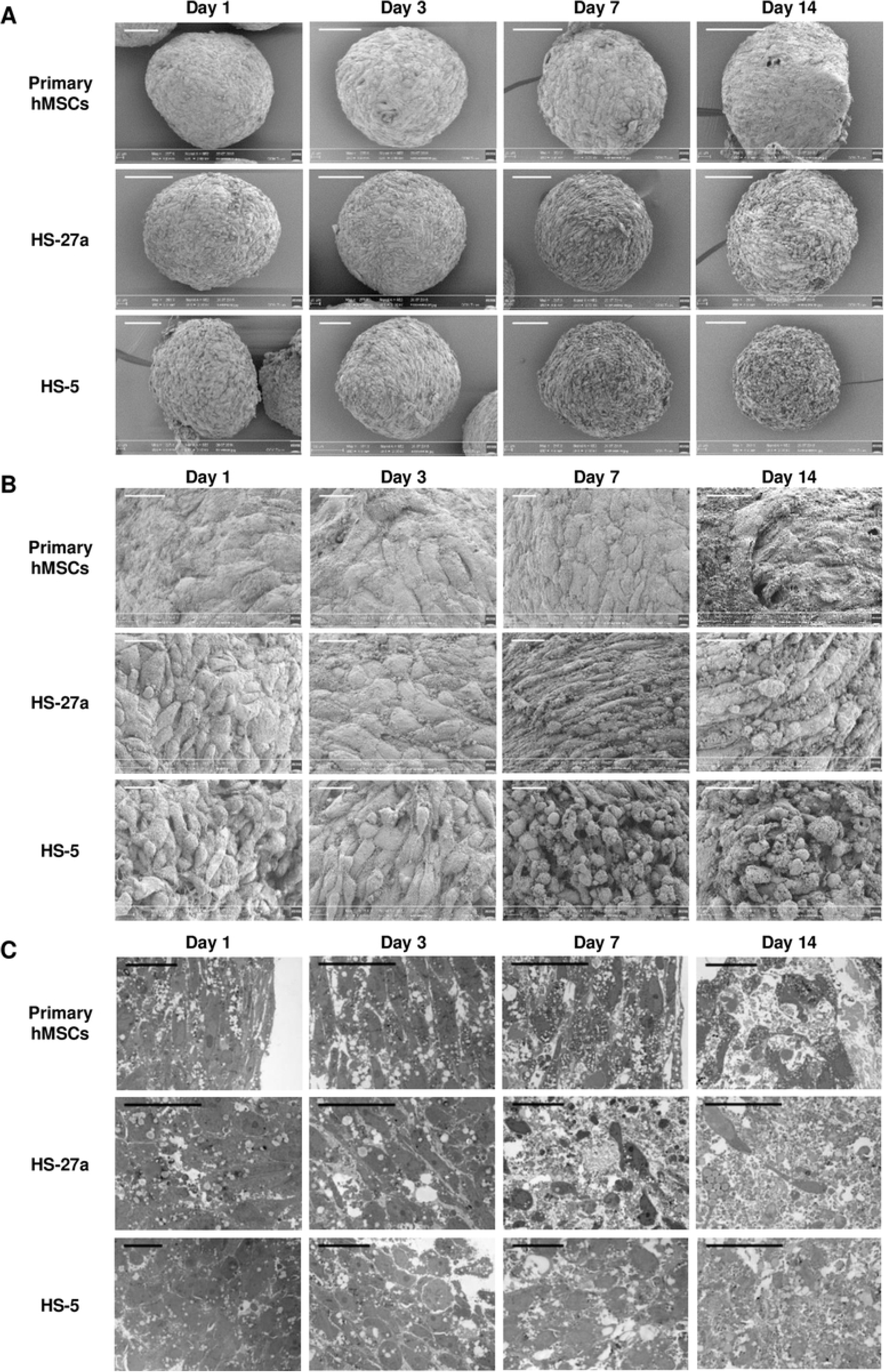
Electron microscopy observation of MSCs-derived spheroids. (A, B) Scanning electron microscopy (SEM) and (C) transmission electronic microscopy (TEM) analysis of spheroids derived from hMSCs, HS-27a and HS-5 cells, over 14 days (scale bars = 100 μm (A), 20 μm (B and C)).

### Cell death and proliferation analyses of the MSCs-derived spheroids

To explain why spheroids had decreased cell number over time, we proposed an imbalance between cell death and cell proliferation. Thus, apoptosis and cell cycle were measured by flow cytometry using 7-AAD/Ki-67 staining (Fig 4A). First, increasing sub-G_0_/G_1_ cell population revealed a strong induction of cell death after 14 days in spheroids obtained with hMSCs, while a more moderate cell death was observed after seven days for the two human cell lines (Fig 4B). Although harvested at the same confluency, primary cells appeared already much more quiescent than HS-27a or HS-5 cells at day 0 (Fig 4C). Then, a significant proportion of cells remained proliferating in spheroids until day 3 for HS-27a and day 7 for HS-5 cells. Remarkably, while closer to HS-27a cells in terms of perimeter and number of cells, MS-5 cells had a massive increase in cell death and almost no proliferation (S1G and S1H Fig). This suggests that, based on proliferation and cell death, the MS-5 cell line is more similar to primary cells than others, probably due to their contact inhibition, which limits their proliferation capacity. Ki-67 detection by immunochemistry, in hMSCs and human cell lines, revealed homogeneous staining at day 1 indicating proliferation in the whole spheroid (Fig 4D) in agreement with a previous study [41]. It also confirmed a lower proliferation rate of hMSCs compared to cell lines and a rapid proliferation arrest with only few Ki-67-positive cells remaining at the periphery of the spheroid at day 3. A progressive decrease in the proliferation for the two human cell lines supported the results obtained by flow cytometry. Interestingly, decreased proliferation appears in the entire spheroid and is not restricted to in-depth localizations. These data showed that spheroids are characterized by imbalance between cell death and proliferation, which may explain the highest loss of cells over time.

**Fig 4.**
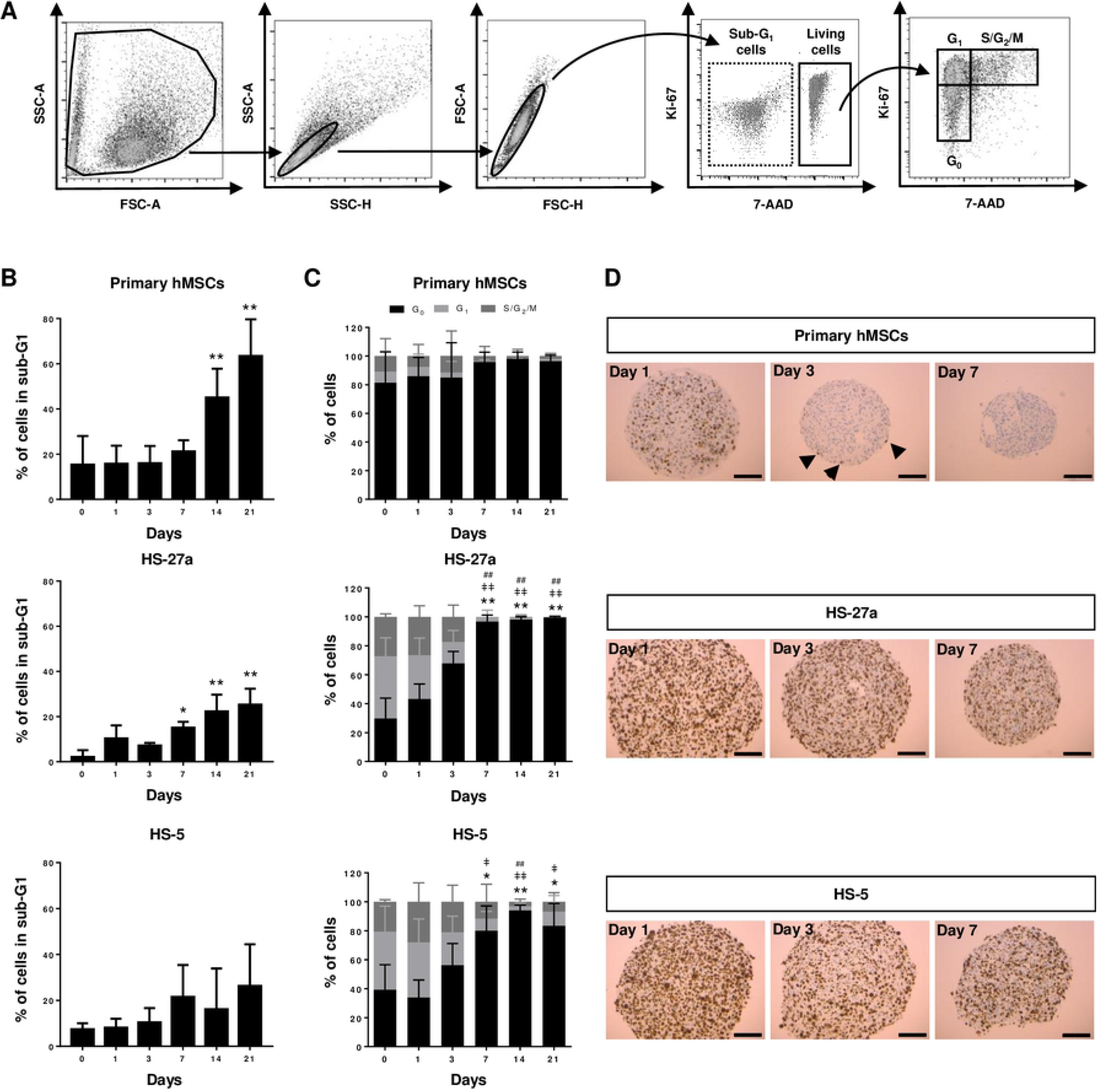
Determination of proliferation and apoptosis of MSCs-derived spheroids. (A-C) Cell cycle analysis of spheroids over 21 days in culture. (A) Representative gating strategy from hMSCs at day 0, (B) sub-G_1_ apoptosis quantification (hMSCs n = 6; HS-27a and HS-5 n = 3) and (C) cell cycle quantification (hMSCs n = 6; HS-27a and HS-5 n = 5; * for G_0_; ǂ for G_1;_ # for S/G_2_/M) (data are mean ± SD; */ǂ/# compared to day 0; */ǂ p ≤ 0.05; **/ǂǂ/## p ≤ 0.01). (D) Immunohistochemistry of Ki-67 at days 1, 3 and 7 for hMSCs-, HS-27a- and HS-5-derived spheroids (scale bars = 100 μm). Arrows indicate Ki-67-positive cells.

### Hypoxia and oxidative stress in MSCs-derived spheroids

Like in tumor spheres [42,43], the appearance of an oxygen gradient and hypoxia in MSCs-derived spheroids [44] has been demonstrated. Carbonic anhydrase IX (CA-IX), a mediator of hypoxia-induced stress response, is commonly used as marker in tumors [45]. Increased CA-IX has been observed in MSCs-derived spheroids, particularly in HS-27a cells (Fig 5A). The pro-survival adaptation to hypoxia occurs mainly through the stabilization of the hypoxia-inducible factors (HIFs). HIFs are key regulators of multiple cell processes, including cell cycle, metabolism, pH control and autophagy. Increasing expression of HIF-1α protein expression has been observed in spheroids over the time, as well as at the mRNA level mainly in hMSCs (Fig 5B). Finally, we examined the expression of *VEGFA*, a standard HIF transcriptionally regulated gene [46]. Its expression in hMSCs- and HS-27a-derived spheroids was already elevated at day 1, but strongly increased at both protein and mRNA levels over time (Fig 5C).

**Fig 5.**
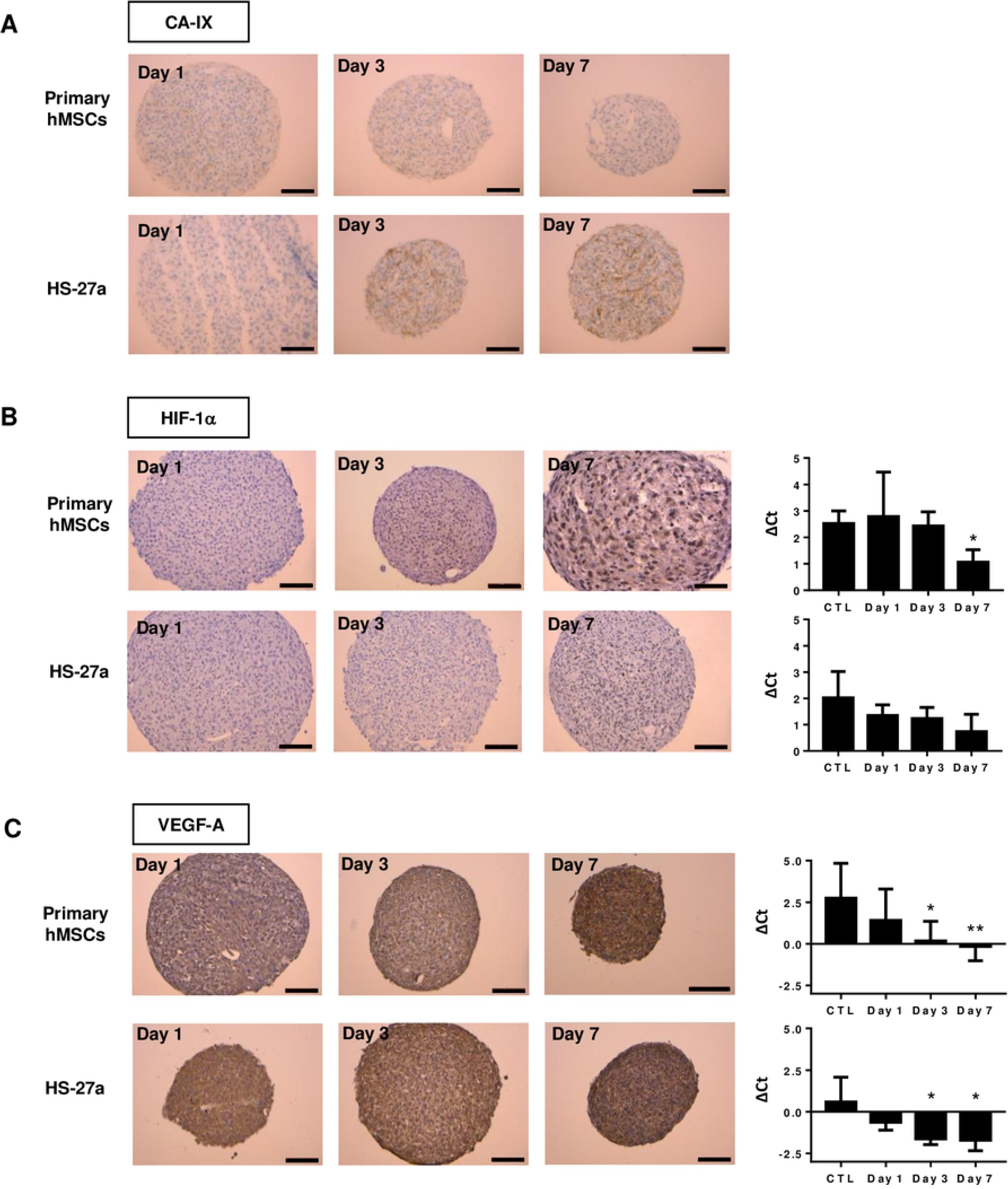
Hypoxia detection of hMSCs- and HS-27a-derived spheroids over 7 days in culture. (A) Immunohistochemistry of CA-IX. (B) Immunohistochemistry and mRNA of HIF-1α. (X) Immunohistochemistry and mRNA expression of VEGF-A. (hMSCs n = 5; HS-27a n = 3; * p ≤ 0.05; ** p ≤ 0.01; scale bars = 100 μm).

In certain circumstances, very low level of oxygen (anoxia) or long exposure to hypoxia may provoke DNA damage and oxidative stress that trigger apoptosis [42]. Besides hypoxia appearance in spheroids, cell aggregation may also stress the cells by itself and increase reactive oxygen species (ROS). Heme oxygenase 1 (HO-1) is induced by a variety of stressors, and is therefore a marker of hypoxia and oxidative stress [47]. Indeed, oxidative stress triggers nuclear relocation of NRF-2, a HO-1 transcription factor, which then leads to antioxidant response through induced expression of antioxidants by HO-1. In the spheroids, we observed a high expression of HO-1 at day 1, which increased over time (Fig 6A). Conversely, among the 24 antioxidant genes (Patent WO2016083742), we found a total of seven genes upregulated in spheroids from the hMSCs and the HS-27a cell line (Fig 6B). Remarkably, of these genes, four (*GPX1*, *PRDX2*, *SOD1* and *SOD2*) were commonly upregulated in both cell types irrespective of their initial expression level.

**Fig 6.**
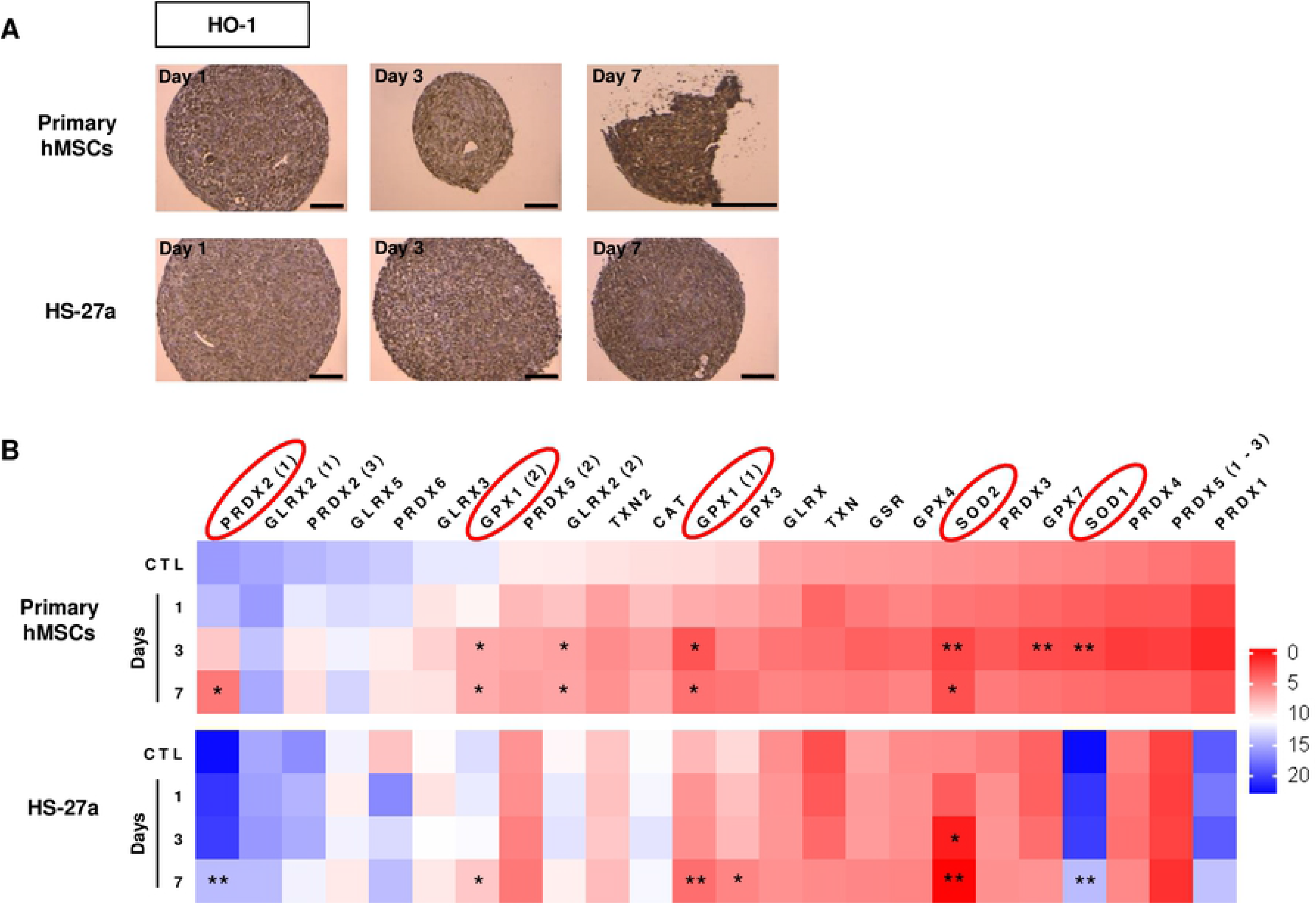
Oxidative stress detection of hMSCs- and HS-27a-derived spheroids over 7 days in culture. (A) Immunohistochemistry of HO-1 (scale bars = 100 μm). (B) Expression of antioxidant genes (n = 3; data are mean; * compared to 2D control (CTL); * p ≤ 0.05; ** p ≤ 0.01).

Together, these data indicate concomitant appearance of hypoxia and oxidative stress in established MSCs-derived spheroids, which could therefore explain initial cell cycle arrest and further apoptosis in prolonged hypoxia [48].

### Stemness in MSCs-derived spheroids

The 2D culture of MSCs critically leads to rapid loss of their pluripotency and supportive functions. In contrast, MSCs-derived spheroids have the potential to maintain stemness that could be demonstrated by the expression of three classical embryonic markers, OCT-4, SOX-2 and NANOG [32]. Furthermore, it has been described that hypoxia transcriptionally regulates these factors in a HIFs-dependent manner [49]. Therefore, in order to validate whether HS-27a behave similarly to hMSCs, we examined the expression of the genes coding for the three factors, over time. Results showed that hMSCs formation was accompanied by upregulation of *OCT4* and *SOX2*, in agreement with previous studies, but surprisingly showed no upregulation of *NANOG* (Fig 7A). HS-27a had similar expression level of the three genes to hMSCs in 2D culture and had progressive increased expression of all three markers (Fig 7B). These data confirmed that, like hMSCs, HS-27a had preserved a stemness capacity that could also be (re)activated during spheroid formation.

**Fig 7.**
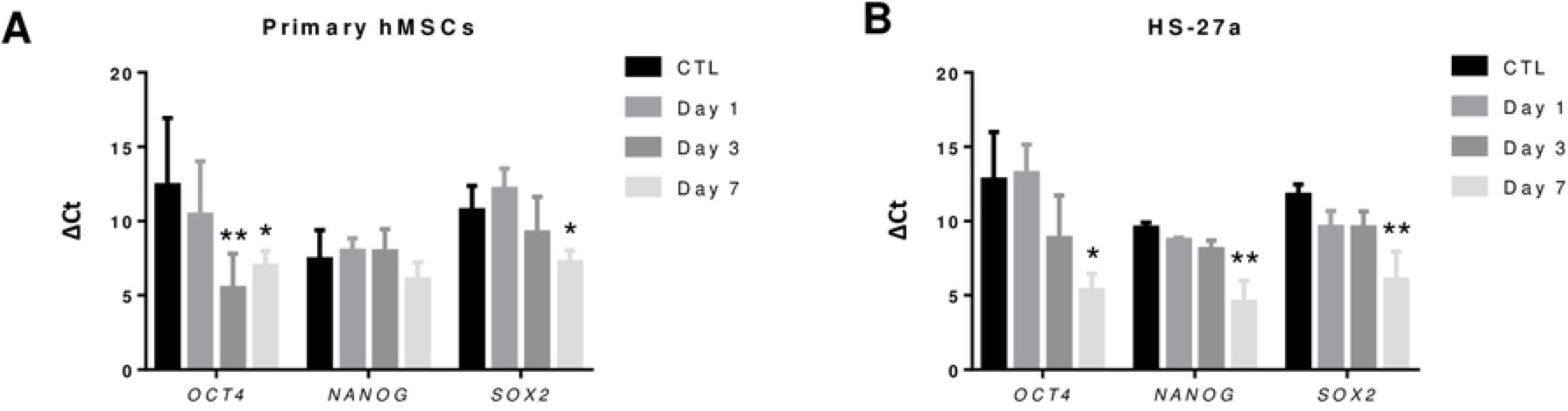
Stemness detection of hMSCs- and HS-27a-derived spheroids over 7 days in culture. (A and B) Gene expression of *OCT4*, *NANOG* and *SOX2* for (A) hMSCs- and (B) HS-27a-derived spheroids (hMSCs n = 5; HS-27a n = 3; * p ≤ 0.05; ** p ≤ 0.01).

## Discussion

In the last decade, studies have shown that MSC spheroids could be a promising model for *in vitro* culture. Indeed, some have demonstrated their benefits in studying cardiac ischemia [50], cerebral ischemia [51], hindlimb ischemia [52] or bone repair [53]. In addition, spheroids may be a good model to study the interaction of normal [17–19,21,22] or malignant hematopoietic cells [20,23] with their microenvironment. For instance, spheroids could be used to study the mechanisms triggering chemoresistance in leukemias [20,23]. However, studies might be limited by the availability of primary human MSCs and the reproducibility due to the different sources, while 2D co-cultures have been for a long time established with cell lines, mostly murine, such as MS-5 or M2-10B4 [11]. In this study, we chose the HS-27a and HS-5 cell lines for their human origin and their capacity to sustain hematopoiesis in co-culture (Roecklein & Torok-Storb, 1995). Nonetheless, in contrast to the murine MS-5 cell line, they do not retain contact inhibition that certainly, although of human origin, have limited their use for long-term culture. We found that both human and murine cell lines, independently of their contact inhibition capacity, were able to provide quick and reproducible spheroids using standard methylcellulose, similarly to hMSCs, with the advantage of keeping the same size over time. The delay to achieve a complete spheroid, 5 h *versus* 10 h for hMSCs and cell lines, respectively, could certainly be attributed to sedimentation speed. In fact, cell lines are much smaller than primaries that could hence sediment faster. On the other hand, this phenomenon might also be attributed to spheroid condensation that could depend on ECM composition. Indeed, hMSCs-derived spheroids appeared more cohesive by SEM.

ECM may also explain, at least partially, shrinking of hMSC-derived spheroids. Shrinking has been previously reported for hMSCs [31,32,35,37,54–56] and has been attributed to induced autophagy [32]. Therefore, we could hypothesize that transformed cell lines may have lower autophagy, which is often induced in reduced or arrest cell growth [57]. Indeed, HS-27a and HS-5 cell lines continue to proliferate until 7 days, unlike hMSCs, and could block autophagy and compensate cell death. However, the number of viable HS-27a decreased over time and no apoptosis has been detected for any of the MSCs before seven days, which could not explain the loss of cells. In agreement, others studies have also demonstrated an induction of apoptosis only after several days [35,54], but not at short term [58].

Studies have already reported oxygen gradients in tumor-spheres [42,43] as well as in MSCs-derived spheroids [44]. The hypoxia response mainly happens through the stabilization of hypoxia-inducible factors (HIFs), which are regulators of multiple biological processes, such as angiogenesis or energetic metabolism. HIFs have an essential pro-survival role by promoting genes, such as those involved in metabolism and autophagy [46]. However, acute and prolonged hypoxia may also trigger cell death through blocking DNA replication and induced oxidative stress [42]. Interestingly, cell lines showed increased hypoxia markers over time, and concomitant decreased cell cycle prior induced apoptosis. This is consistent with induced oxidative stress revealed by increased expression of HO-1 and antioxidant response.

## Conclusions

Overall these data indicate that, like hMSCs, MSC cell lines make reproductible and easily handled spheroids. Remarkably, the HS-27a cell line more closely resemble primary cells than the HS-5 line. This is of a particular interest, since HS-27a has been shown to provide better support to HSCs [38–40]. Thus, this model could help in understanding mechanisms involved in MSC physiology and may be a simple model to study cell interactions in the hematopoietic niche. The model could also be extended to research metastatic process as previously described for breast cancer [28].

## Supporting information

**S1 Fig. Spheroids formation of mouse MS-5 cell line.** (A) Microscopy analysis over 21 days in culture (scale bars = 100 μm). (B) Perimeter was measured with an arbitrary unit; each experiment is the mean of at least 10 spheroids (n = 3; data are mean ± SD). (C) Number of living cells per spheroids over 21 days in culture (n = 3; each experiment is the mean of 12 spheroids). (D, E) Scanning electron microscopy (SEM) and (F) transmission electronic microscopy (TEM) analysis over 14 days (scale bars = 100 μm (D), 20 μm (E and F). (G) Sub-G_1_ apoptosis quantification (n = 3) and (H) cell cycle quantification over 21 days in culture (n = 3; data are mean ± SD).

**S1 Video.** A representative time-lapse video of spheroid formation. 30 000 primary MSCs seeded into U-bottomed 96-well, in medium containing 0.5 % of methylcellulose (Methocult™ SF H4236) were followed via a Nikon Eclipse TI-S microscope for 24 hours.

**S2 Video.** A representative time-lapse video of spheroid formation. 30 000 HS-27a cells seeded into U-bottomed 96-well, in medium containing 0.5 % of methylcellulose (Methocult™ SF H4236) were followed via a Nikon Eclipse TI-S microscope for 24 hours.

**S3 Video.** A representative time-lapse video of spheroid formation. 30 000 HS-5 cells seeded into U-bottomed 96-well, in medium containing 0.5 % of methylcellulose (Methocult™ SF H4236) were followed via a Nikon Eclipse TI-S microscope for 24 hours.

**S4 Video.** A representative time-lapse video of spheroid formation. 30 000 MS-5 cells seeded into U-bottomed 96-well, in medium containing 0.5 % of methylcellulose (Methocult™ SF H4236) were followed via a Nikon Eclipse TI-S microscope for 24 hours.

## References

1. Domenech J. What Are Mesenchymal Stromal Cells? Origin and Discovery of Mesenchymal Stromal Cells. In: Mesenchymal Stromal Cells as Tumor Stromal Modulators. 2017. p. 1–37.

2. Dominici M, Le Blanc K, Mueller I, Slaper-Cortenbach I, Marini F, Krause D, et al. Minimal criteria for defining multipotent mesenchymal stromal cells. The International Society for Cellular Therapy position statement. Cytotherapy. 2006;8(4):315–7.

3. Makino S, Fukuda K, Miyoshi S, Konishi F, Kodama H, Pan J, et al. Cardiomyocytes can be generated from marrow stromal cells in vitro. J Clin Invest. 1999;103(5):697–705.

4. Sanchez-Ramos J, Song S, Cardozo-Pelaez F, Hazzi C, Stedeford T, Willing A, et al. Adult bone marrow stromal cells differentiate into neural cells in vitro. Exp Neurol. 2000;164(2):247–56.

5. Spees JL, Olson SD, Ylostalo J, Lynch PJ, Smith J, Perry A, et al. Differentiation, cell fusion, and nuclear fusion during ex vivo repair of epithelium by human adult stem cells from bone marrow stroma. Proc Natl Acad Sci. 2003;100(5):2397–402.

6. Hong SH, Gang EJ, Jeong JA, Ahn C, Hwang SH, Yang IH, et al. In vitro differentiation of human umbilical cord blood-derived mesenchymal stem cells into hepatocyte-like cells. Biochem Biophys Res Commun. 2005;330(4):1153–61.

7. Yan L, Zheng D, Xu RH. Critical role of tumor necrosis factor signaling in mesenchymal stem cell-based therapy for autoimmune and inflammatory diseases. Front Immunol. 2018;9:1658.

8. Su P, Tian Y, Yang C, Ma X, Wang X, Pei J, et al. Mesenchymal stem cell migration during bone formation and bone diseases therapy. Int J Mol Sci. 2018;19(8):E2343.

9. Perez JR, Kouroupis D, Li DJ, Best TM, Kaplan L, Correa D. Tissue Engineering and Cell-Based Therapies for Fractures and Bone Defects. Front Bioeng Biotechnol. 2018;6(105):1–23.

10. Pinho S, Frenette PS. Haematopoietic stem cell activity and interactions with the niche. Nat Rev Mol Cell Biol. 2019;20(5):303–20.

11. Vaidya A, Kale V. Hematopoietic stem cells, their niche, and the concept of co-culture systems: A critical review. J Stem Cells. 2015;10(1):13–31.

12. Dhami SPS, Kappala SS, Thompson A, Szegezdi E. Three-dimensional ex vivo co-culture models of the leukaemic bone marrow niche for functional drug testing. Drug Discov Today. 2016;21(9):1464–71.

13. Cesarz Z, Tamama K. Spheroid Culture of Mesenchymal Stem Cells. Stem Cells Int. 2016;2016:e9176357.

14. Lin RZ, Chang HY. Recent advances in three-dimensional multicellular spheroid culture for biomedical research. Biotechnol J. 2008;3(9–10):1172–84.

15. Baraniak PR, McDevitt TC. Scaffold-free culture of mesenchymal stem cell spheroids in suspension preserves multilineage potential. Cell Tissue Res. 2012;347(3):701–11.

16. Ghazanfari R, Li H, Zacharaki D, Lim HC, Scheding S. Human Non-Hematopoietic CD271pos/CD140alow/neg Bone Marrow Stroma Cells Fulfill Stringent Stem Cell Criteria in Serial Transplantations. Stem Cells Dev. 2016;25(21):1652–8.

17. Futrega K, Atkinson K, Lott WB, Doran MR. Spheroid Coculture of Hematopoietic Stem/Progenitor Cells and Monolayer Expanded Mesenchymal Stem/Stromal Cells in Polydimethylsiloxane Microwells Modestly Improves In Vitro Hematopoietic Stem/Progenitor Cell Expansion. Tissue Eng Part C Methods. 2017;23(4):200–18.

18. De Barros APDN, Takiya CM, Garzoni LR, Leal-Ferreira ML, Dutra HS, Chiarini LB, et al. Osteoblasts and bone marrow mesenchymal stromal cells control hematopoietic stem cell migration and proliferation in 3D in vitro model. PLoS One. 2010;5(2):e9093.

19. Isern J, Martín-Antonio B, Ghazanfari R, Martín AM, López JA, DelToro R, et al. Self-Renewing Human Bone Marrow Mesenspheres Promote Hematopoietic Stem Cell Expansion. Cell Rep. 2013;3(5):1714–24.

20. Bruce A, Evans R, Mezan R, Shi L, Moses BS, Martin KH, et al. Three-dimensional microfluidic tri-culture model of the bone marrow microenvironment for study of acute lymphoblastic leukemia. PLoS One. 2015;10(10):e0140506.

21. Wuchter P, Saffrich R, Giselbrecht S, Nies C, Lorig H, Kolb S, et al. Microcavity arrays as an in vitro model system of the bone marrow niche for hematopoietic stem cells. Cell Tissue Res. 2016;364(3):573–84.

22. Leisten I, Kramann R, Ventura Ferreira MS, Bovi M, Neuss S, Ziegler P, et al. 3D co-culture of hematopoietic stem and progenitor cells and mesenchymal stem cells in collagen scaffolds as a model of the hematopoietic niche. Biomaterials. 2012;33(6):1736–47.

23. Aljitawi OS, Li D, Xiao Y, Zhang D, Ramachandran K, Stehno-Bittel L, et al. A novel three-dimensional stromal-based model for in vitro chemotherapy sensitivity testing of leukemia cells. Leuk Lymphoma. 2014;55(2):378–91.

24. Sart S, Tsai A-C, Li Y, Ma T. Three-dimensional aggregates of mesenchymal stem cells: cellular mechanisms, biological properties, and applications. Tissue Eng Part B Rev. 2014;20(5):365–80.

25. Egger D, Tripisciano C, Weber V, Dominici M, Kasper C. Dynamic Cultivation of Mesenchymal Stem Cell Aggregates. Bioengineering. 2018;5(2):1–15.

26. Ong SM, Zhang C, Toh YC, Kim SH, Foo HL, Tan CH, et al. A gel-free 3D microfluidic cell culture system. Biomaterials. 2008;29(22):3237–44.

27. Saleh FA, Frith JE, Lee JA, Genever PG. Three-Dimensional In Vitro Culture Techniques for Mesenchymal Stem Cells. In: Progenitor Cells: Methods and Protocols. 2012. p. 31–45.

28. Cavnar SP, Rickelmann AD, Meguiar KF, Xiao A, Dosch J, Leung BM, et al. Modeling Selective Elimination of Quiescent Cancer Cells from Bone Marrow. Neoplasia. 2015;17(8):625–33.

29. Itoh K, Tezuka H, Sakoda H, Konno M, Nagata K, Uchiyama T, et al. Reproducible establishment of hemopoetic supportive stromal cell lines from murine bone marrow. Exp Hematol. 1989;17(2):145–53.

30. Saleh FA, Whyte M, Genever PG. Effects of endothelial cells on human mesenchymal stem cell activity in a three-dimensional in vitro model. Eur Cells Mater. 2011;22:242–57.

31. Shearier E, Xing Q, Qian Z, Zhao F. Physiologically Low Oxygen Enhances Biomolecule Production and Stemness of Mesenchymal Stem Cell Spheroids. Tissue Eng Part C. 2016;22(4):360–9.

32. Pennock R, Bray E, Pryor P, James S, McKeegan P, Sturmey R, et al. Human cell dedifferentiation in mesenchymal condensates through controlled autophagy. Sci Rep. 2015;5:e13113.

33. Matak D, Brodaczewska KK, Lipiec M, Szymanski Ł, Szczylik C, Czarnecka AM. Colony, hanging drop, and methylcellulose three dimensional hypoxic growth optimization of renal cell carcinoma cell lines. Cytotechnology. 2017;69(4):565–78.

34. Foty R. A Simple Hanging Drop Cell Culture Protocol for Generation of 3D Spheroids. J Vis Exp. 2011;20(51):1–4.

35. Schmal O, Seifert J, Schäffer T, Walter CB, Aicher WK, Klein G. Hematopoietic Stem and Progenitor Cell Expansion in Contact with Mesenchymal Stromal Cells in a Hanging Drop Model Uncovers Disadvantages of 3D. Stem Cells Int. 2015;2016:e4148093.

36. Blocki A, Wang Y, Koch M, Peh P, Beyer S, Law P, et al. Not All MSCs Can Act as Pericytes : Functional In Vitro Assays to Distinguish Pericytes from Other Mesenchymal Stem Cells in Angiogenesis. Stem Cells Dev. 2013;22(17):2347–55.

37. Redondo-Castro E, Cunningham CJ, Miller J, Cain S, Allan SM, Pinteaux E. Generation of Human Mesenchymal Stem Cell 3D Spheroids Using Low-binding Plates. Bio Protoc. 2018;8(16):e2968.

38. Roecklein BA, Torok-Storb B. Functionally distinct human marrow stromal cell lines immortalized by transduction with the human papilloma virus E6/E7 genes. Blood. 1995;85(4):997–1005.

39. Torok-Storb B, Iwata M, Graf L, Gianotti J, Horton H, Byrne MC. Dissecting the marrow microenvironment. In: Annals of the New York Academy of Sciences. 1999. p. 164–70.

40. Iwata M, Sandstrom RS, Delrow JJ, Stamatoyannopoulos JA, Torok-Storb B. Functionally and phenotypically distinct subpopulations of marrow stromal cells are fibroblast in origin and induce different fates in peripheral blood monocytes. Stem Cells Dev. 2014;23(7):729–40.

41. Li Y, Guo G, Li L, Chen F, Bao J, Shi Y jun, et al. Three-dimensional spheroid culture of human umbilical cord mesenchymal stem cells promotes cell yield and stemness maintenance. Cell Tissue Res. 2015;360(2):297–307.

42. Riffle S, Hegde RS. Modeling tumor cell adaptations to hypoxia in multicellular tumor spheroids. J Exp Clin Cancer Res. 2017;36(1):e102.

43. Leek R, Grimes DR, Harris AL, McIntyre A. Methods: Using Three-Dimensional Culture (Spheroids) as an In Vitro Model of Tumour Hypoxia. In: Tumor microenvironment. 2016. p. 167–96.

44. Sharma MB, Limaye LS, Kale VP. Mimicking the functional hematopoietic stem cell niche in vitro: Recapitulation of marrow physiology by hydrogel-based three-dimensional cultures of mesenchymal stromal cells. Haematologica. 2012;97(5):651–60.

45. McDonald PC, Dedhar S. Carbonic anhydrase IX (CAIX) as a mediator of hypoxia-induced stress response in cancer cells. In: SubCellular Biochemistry. 2014. p. 255–69.

46. Zhang CC, Sadek HA. Hypoxia and Metabolic Properties of Hematopoietic Stem Cells. Antioxid Redox Signal. 2014;20(12):1891–901.

47. Dunn LL, Midwinter RG, Ni J, Hamid HA, Parish CR, Stocker R. New insights into intracellular locations and functions of heme oxygenase-1. Antioxidants Redox Signal. 2014;20(11):1723–42.

48. Riffle S, Pandey RN, Albert M, Hegde RS. Linking hypoxia, DNA damage and proliferation in multicellular tumor spheroids. BMC Cancer. 2017;17(1):1–12.

49. Drela K, Sarnowska A, Siedlecka P, Szablowska-Gadomska I, Wielgos M, Jurga M, et al. Low oxygen atmosphere facilitates proliferation and maintains undifferentiated state of umbilical cord mesenchymal stem cells in an hypoxia inducible factor-dependent manner. Cytotherapy. 2014;16(7):881–92.

50. Ceccaldi C, Bushkalova R, Alfarano C, Lairez O, Calise D, Bourin P, et al. Evaluation of polyelectrolyte complex-based scaffolds for mesenchymal stem cell therapy in cardiac ischemia treatment. Acta Biomater. 2014;10(2):901–11.

51. Guo L, Ge J, Zhou Y, Wang S, Zhao RCH, Wu Y. Three-Dimensional Spheroid-Cultured Mesenchymal Stem Cells Devoid of Embolism Attenuate Brain Stroke Injury After Intra-Arterial Injection. Stem Cells Dev. 2013;23(9):978–89.

52. Bhang SH, Lee S, Shin J-Y, Lee T-J, Kim B-S. Transplantation of Cord Blood Mesenchymal Stem Cells as Spheroids Enhances Vascularization. Tissue Eng Part A. 2012;18(19–20):2138–47.

53. Yamaguchi Y, Ohno J, Sato A, Kido H, Fukushima T. Mesenchymal stem cell spheroids exhibit enhanced in-vitro and in-vivo osteoregenerative potential. BMC Biotechnol. 2014;14(105):1–10.

54. Tsai A-C, Liu Y, Yuan X, Ma T. Compaction, fusion, and functional activation of three-dimensional human mesenchymal stem cell aggregate. Tissue Eng Part A. 2015;21(9–10):1705–19.

55. Kim M, Yun H, Young D, Byung P, Choi H. Three-Dimensional Spheroid Culture Increases Exosome Secretion from Mesenchymal Stem Cells. Tissue Eng Regen Med. 2018;15(4):427–36.

56. Bellotti C, Duchi S, Bevilacqua A, Lucarelli E, Piccinini F. Long term morphological characterization of mesenchymal stromal cells 3D spheroids built with a rapid method based on entry-level equipment. Cytotechnology. 2016;1–12.

57. Neufeld TP. Autophagy and cell growth - the yin and yang of nutrient responses. J Cell Sci. 2012;125(10):2359–68.

58. Ho SS, Hung BP, Heyrani N, Lee MA, Leach JK. Hypoxic Preconditioning of Mesenchymal Stem Cells with Subsequent Spheroid Formation Accelerates Repair of Segmental Bone Defects. Stem Cells. 2018;36(9):1393–403.

